# Enhancing Radiation-induced Reactive Oxygen Species Generation Through Mitochondrial Transplantation in Human Glioblastoma

**DOI:** 10.1101/2024.10.20.619301

**Authors:** Kent L Marshall, Murugesan Velayutham, Valery V. Khramtsov, Alan Mizener, Christopher P Cifarelli

**Affiliations:** Department of Neurosurgery, Rockefeller Neuroscience Institute, West Virginia University, Morgantown, WV, USA; Department of Biochemistry and Molecular Medicine, West Virginia University, Morgantown, WV, USA; West Virginia University Cancer Institute, Morgantown, WV, USA; Department of Radiation Oncology, West Virginia University, Morgantown, WV, USA

**Keywords:** glioblastoma, mitochondria, ROS, EPR, radiation, cell-penetrating peptide, RBE

## Abstract

Glioblastoma (GBM) is the most aggressive primary brain malignancy in adults, with high recurrence rates and resistance to standard therapies. This study explores mitochondrial transplantation as a novel method to enhance the radiobiological effect (RBE) of ionizing radiation (IR) by increasing mitochondrial density in GBM cells, potentially boosting reactive oxygen species (ROS) production and promoting radiation-induced cell death. Using cell-penetrating peptides (CPPs), mitochondria were transplanted into GBM cell lines U3020 and U3035. Transplanted mitochondria were successfully incorporated into recipient cells, increasing mitochondrial density significantly. Mitochondrial chimeric cells demonstrated enhanced ROS generation post-irradiation, as evidenced by increased electron paramagnetic resonance (EPR) signal intensity and fluorescent ROS assays. The transplanted mitochondria retained functionality and viability for up to 14 days, with mitochondrial DNA (mtDNA) sequencing confirming high transfection and retention rates. Notably, mitochondrial transplantation was feasible in radiation-resistant GBM cells, suggesting potential clinical applicability. These findings support mitochondrial transplantation as a promising strategy to overcome therapeutic resistance in GBM by amplifying ROS-mediated cytotoxicity, warranting further investigation into its efficacy and mechanisms *in vivo*.

## Introduction

Glioblastoma remains the most common, aggressive, and lethal primary brain malignancy in adults despite incremental advances in management over the past three decades [1, 2]). While the standard of care (SOC) for management of GBM typically includes maximal safe surgical resection followed by chemoradiotherapy, disease progression is invariable. In fact, the 2-year local recurrence rate of GBM is estimated to be 90% despite multimodal therapy with remaining malignant cells often progressing with phenotypes which are more resistant to pharmacological intervention and radiotherapy [3–5]. Further complicating the issue, there are currently no consensuses on the SOC for recurrent GBM making the establishment of new options for therapy for the management of both primary and recurrent GBM paramount for improving patient survival and quality of life.

Radiation treatment has played a pivotal role in the treatment of GBM for decades and represents one the few therapies where Level 1 clinical evidence supports use. As with other malignancies, the DNA damage incurred after exposure to ionizing radiation (IR) impacts rapidly dividing GBM cells and leads to local toxic effects in the presence of impaired DNA damage response (DDR) mechanisms [6]. Central to this cytotoxic process is the intracellular production of reactive oxygen species (ROS), primarily by the radiolysis of water and the generation of hydroxyl radicals (OH·), with downstream generation of other radical species, such as superoxide anions (O_2_·) and hydrogen peroxide (H_2_O_2_). In total, all ROS can damage macromolecular structures like lipids, proteins, and DNA [7, 8]). Under normal physiological circumstances, the mitochondria are the primary source of intracellular ROS, designated mtROS [9, 10]. IR can induce OH· through radiolysis of water within the mitochondria; however it can also alter the structure and function of mitochondria leading to breakdown of the electron transport chain (ETC) and oxidative phosphorylation leading to sustained production of ROS [11, 12]. These damaged mitochondria contribute to apoptotic responses directly through expulsion of pro-apoptotic signaling molecules like cytochrome C and caspases [13].

While radiation improves the survival and quality of life in GBM patients, recurrence and associated radiation resistant phenotypes lead to eventual therapeutic failure. There have been recent attempts, to develop novel compounds or repurpose existing pharmaceuticals to sensitize or re-sensitize GBM to radiotherapy including temozolomide, quisinostat, difluorodeoxyuridine, among others [14, 15]. However, these compounds have only demonstrated modest improvement in radiotherapy for GBM and guidelines currently do not support the use of radiosensitizers in CNS disease [16, 17].

Moving beyond direct pharmaceutical-based interventions, precision medicine efforts could be focused on improving the radiobiological effect (RBE) of IR through increasing the density of mitochondria within a population of tumors cells targeted for radiation-induced cell death.

Mitochondrial transplantation has been described in context of deficiency diseases such as, myoclonic epilepsy with ragged red fibers (MERRF), mitochondrial encephalomyopathy with lactic acidosis and stroke-like episodes (MELAS), and Leber hereditary optic neuropathy (LHON) [18, 19]. Regardless of the success, or lack thereof, in these models, the current literature supports the successful incorporation of exogenous mitochondria into donor cells using cell-penetrating peptides (CPPs) [20]. Increasing the number of mitochondria in neoplastic cells will potentially allow for the generation of more ROS and pro-apoptotic molecules and lead to an improved response to radiation and radiobiological effect. Therefore, this study aims to provide proof-of-concept for the transfer of exogenous mitochondria into GBM cell lines in order to evaluate the feasibility, efficacy, and durability of mitochondrial transplantation as a precision medicine tool for radiotherapy.

## Results

### Exogenous allogenic and autologous transplant is possible between discrete GBM cell lines

Mitochondria extracted from donor U3035 cells were successfully incorporated into the cytoplasm of recipient U3035 cells at a concentration of 100µg of mitochondrial protein/5.0×10^5^ U3035 cells, as a proof of concept, and signals persisted through post-transplant day 7 (**Figure 1**). Mitochondria extracted from donor U3020 cells were also successfully incorporated into the cytoplasm of recipient U3035 cells at a concentration of 100µg of mitochondrial protein/5.0×10^5^ U3035 cells and signals persisted through post-transplant day 7 (**Figure 1**) generating allogenic U3035Mt20-chimeras. Increasing the concentration of donor U3020 mitochondria to 300µg of mitochondrial protein/5.0×10^5^ recipient U3035 cells increased the fluorescent signal of resultant mitochondrial chimeric cells by 6.32-fold as compared to the measured at post-transplant day 7, and generated U3035Mt20-3X chimeras (p=0.002) (**Figure 2**). Autologous, allogenic, and 3X allogenic mitochondrial chimeras remained viable for 30 days post-transplant. However, vital dye signals were lost following visualization at post-transplant day 14.

**Figure 1.**
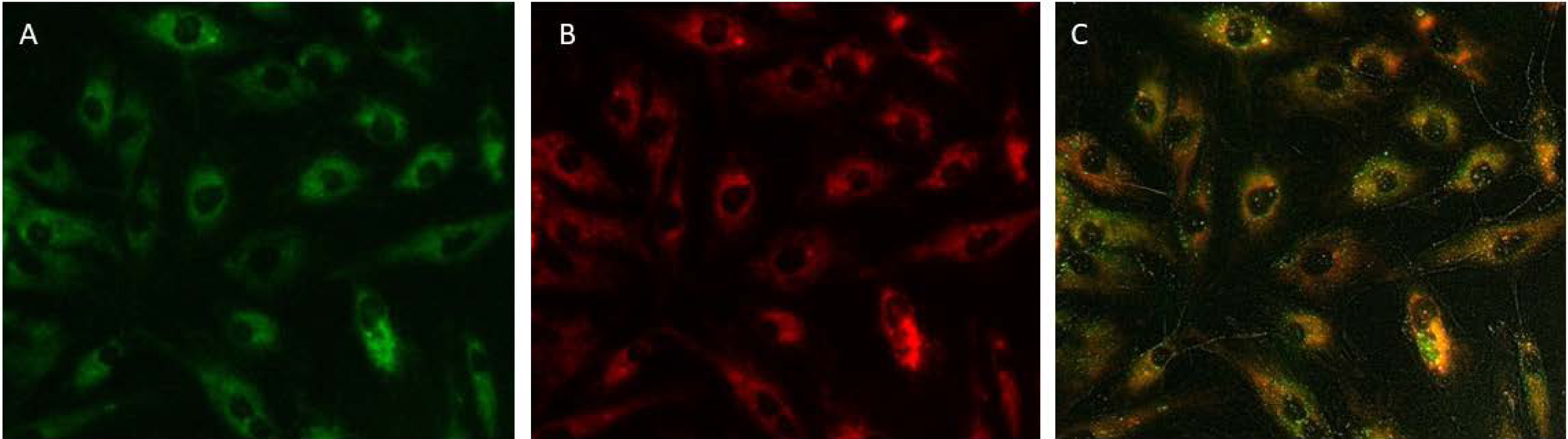
Proof of Concept for Allogenic Mitochondrial Transplantation. Representative images of **A).** Native mitochondria in U3035 cells stained with 200nM MitoTracker® Green vital dye. **B).** Transplanted mitochondria from U3020 cells stained with 200nM MitoTracker® Red vital dye. **C)**. Overlay image of the green and red fluorescent channels highlighting successful transfer. Images were obtained at post-transplant at 48hrs at 100X magnification.

**Figure 2.**
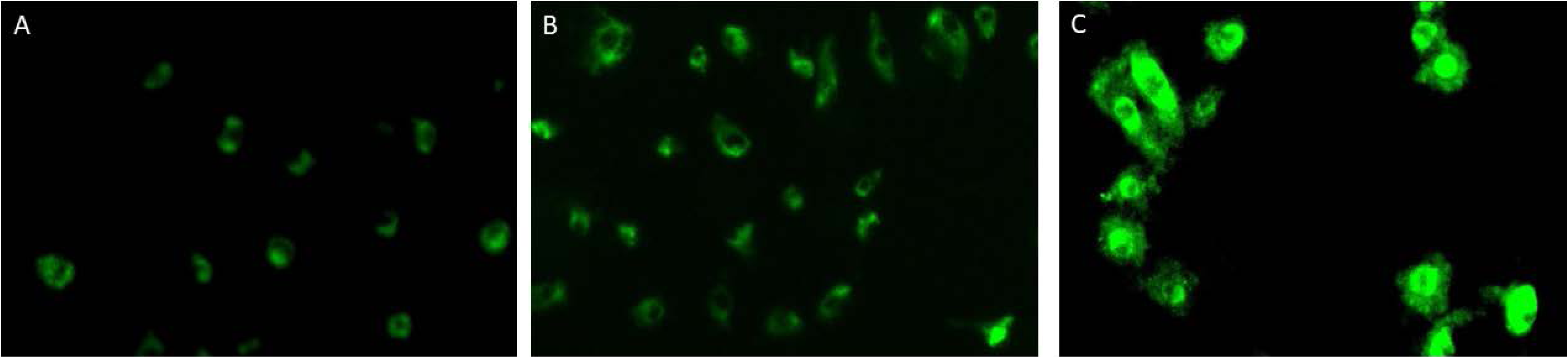
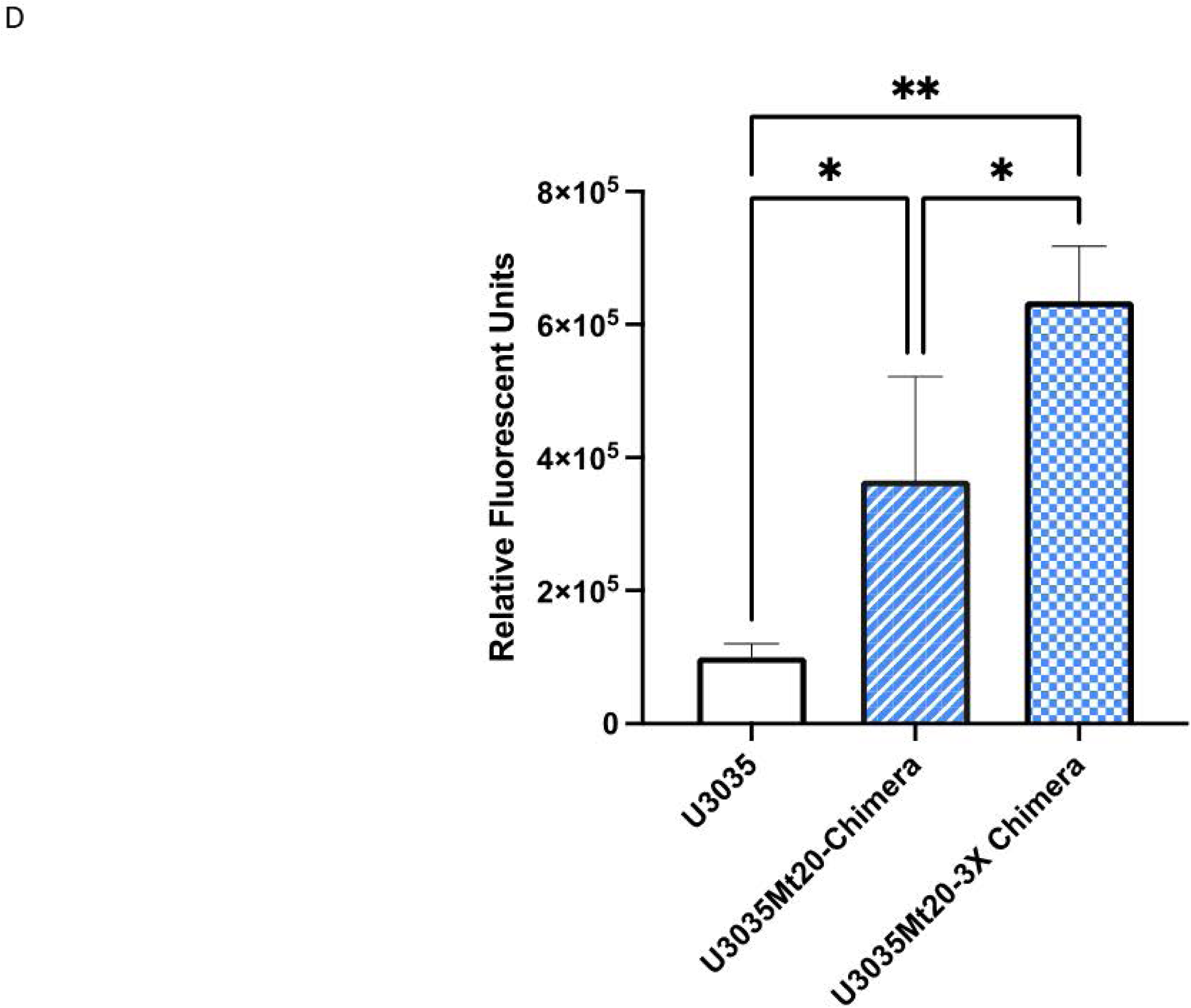
Feasibility of Scaling up Mitochondrial Transplantation. **A)**. Native mitochondria within U3035 cells stained with 200nm MtioTracker® Green vital dye. **B)**. Post-transplant image of allogenic U3020 mitochondria (100µg of mitochondrial protein) stained with 200nm MitoTracker® Green vital dye in U3035 cells with native mitochondria also stained with 200nm MitoTracker® Green vital dye (U3035Mt20 chimera). **C)**. Post-transplant image of allogenic U3020 mitochondria (300µg of mitochondrial protein) stained with 200nm MitoTracker® Green vital dye in U3035 cells with native mitochondria also stained with 200nm MitoTracker® Green vital dye (U3035Mt20-3x chimera). **D)**. Quantification of fluorescent signal using Image J and reported as mean relative fluorescent units ± the standard deviation. Means were: 1.04×10^5^±2.01×10^4^ for U3035, 3.65×10^5^±1.57×10^5^ for U3035Mt20 chimeras, and 6.45×10^5^±8.25×10^4^. One way ANOVA was conducted using a Tukey’s multiple comparisons pos-hoc test to determine significance in the difference of means between groups. All data were analyzed using GraphPad Prism 10 software. *p<0.05 and **p<0.01.

### Mitochondrial density was distinct between individual GBM lines and chimeras

Mitochondrial densities were determined using total nanograms of mitochondrial protein per cell following mitochondrial isolation procedures. U3035 cells had a baseline mean mitochondrial density of 0.135±0.04 ng/cell while U3020 cells had a baseline mean density of 0.28±0.1 ng/cell (**Figure 3**).

**Figure 3.**
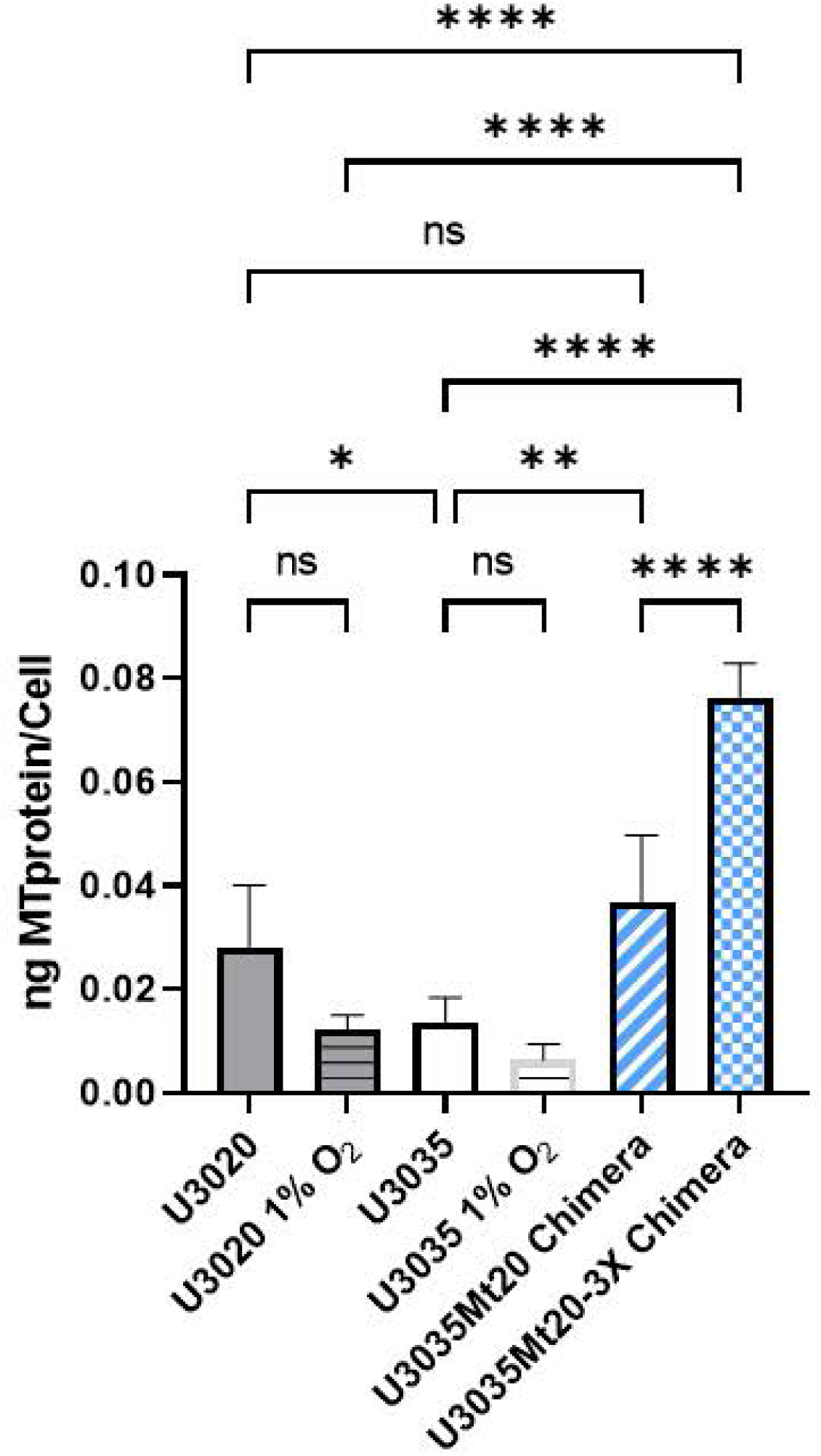
Quantification of Mitochondrial Density. Mitochondrial densities from each cell line were obtained by dividing the total mass of mitochondrial protein (ug) that were extracted from individual lines by the number of live cells following dissociation and mitochondrial isolation. To demonstrate the predictable dynamic nature of mitochondrial density, and intact mitochondrial signaling response in the U3020 and U3035 lines, mitochondrial densities were also included from cells cultured under hypoxic conditions. One-way ANOVA was conducted using a Tukey’s multiple comparisons pos-hoc test to determine significance in the difference of means between groups. All data were analyzed using GraphPad Prism 10 software. * p<0.05 and ** p<0.01.

Following allogenic transplant, U3035Mt20-chimeras had a density of 0.38±0.1 ng/cell 7 days post-transplant, which represented a significant increase over the baseline U3035 donor measurement (p=0.003). Furthermore, the U3035Mt20-3X chimeras demonstrated a mitochondrial density of 0.76±0.07 ng/cell 7 days post-transplant, which represents a significant increase in density when compared to both the U3035 donor cells (p<0.0001) and U3035Mt20-chimeras (p<0.0001). In order to demonstrate the dynamic nature of the mitochondrial density within GBM cells, U3020 and U3035 cells were cultured under hypoxic conditions. U3020 and U3035 hypoxic cells had mitochondrial densities of 0.121±0.025 and 0.063±0.04 ng/cell respectively.

### Mitochondrial Chimeric cells increased the amount of reactive oxygen species generated following exposure to ionizing radiation

When exposed to 8Gy of low-energy X-ray radiation (220kV), U3020 cells and U3035 cells exhibited similar generation of hydroxyl radicals as measured by absolute signal intensity of EPR (0.248±0.006 and 0.246±0.007, respectively) (**Figure 4**). U3035Mt20-chimeras showed a significant increase in EPR signal intensity (0.311±0.01) as compared to the U3035 cells alone (p=0.02). U3035Mt20-3X chimeras showed a significantly higher EPR signal intensity at 0.453±0.03 as compared to both U3035 cells and U3035Mt20-chimeras (p<0.0001 for both comparisons). There were no appreciable differences between allogenic and autologous chimeric cell lines. These data were confirmed using a fluorescent pan-ROS assay (**Figure 5**). U3035Mt20-3X chimeras showed a marked increase in overall ROS following 8Gy irradiation at 0.5 and 24h vs their 0Gy baseline as compared to radiation responses in the native U3035 cells. U3035Mt20-3X chimeras exhibited a fluorescent signal intensity 2.3-fold higher at 0.5h and 3.3-fold higher at 24h. Change in ROS signal intensity was greater in U3035Mt20-3X chimeras when compared to U3035 (p=0.005) but not U3020 (p=0.14) at 0.5h post-irradiation. U3035 and U3020 cells showed a response similar to one another following 8Gy irradiation at 0.5hr with 1.7- and 1.2-fold increases, respectively (p=0.26).

**Figure 4.**
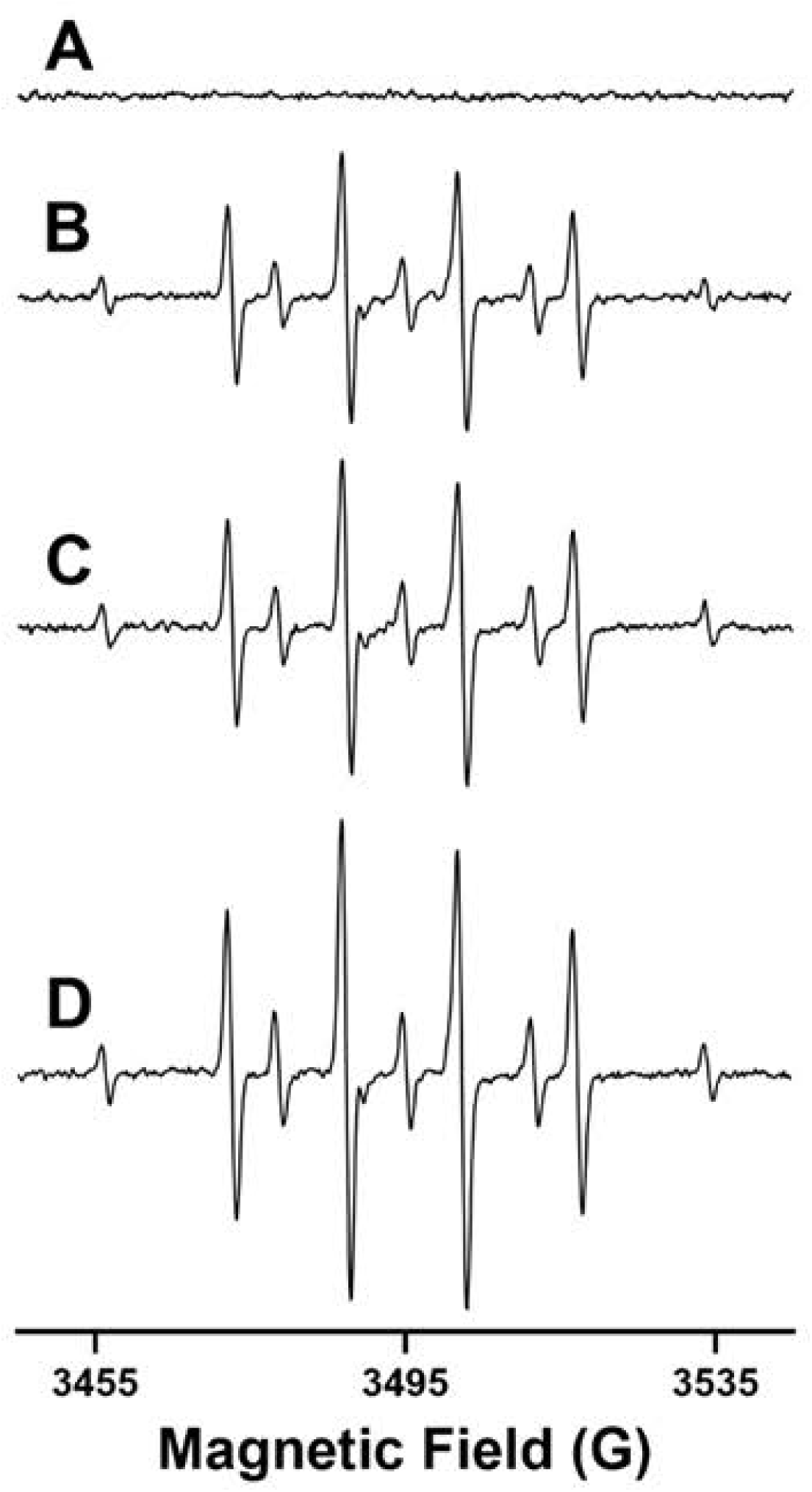
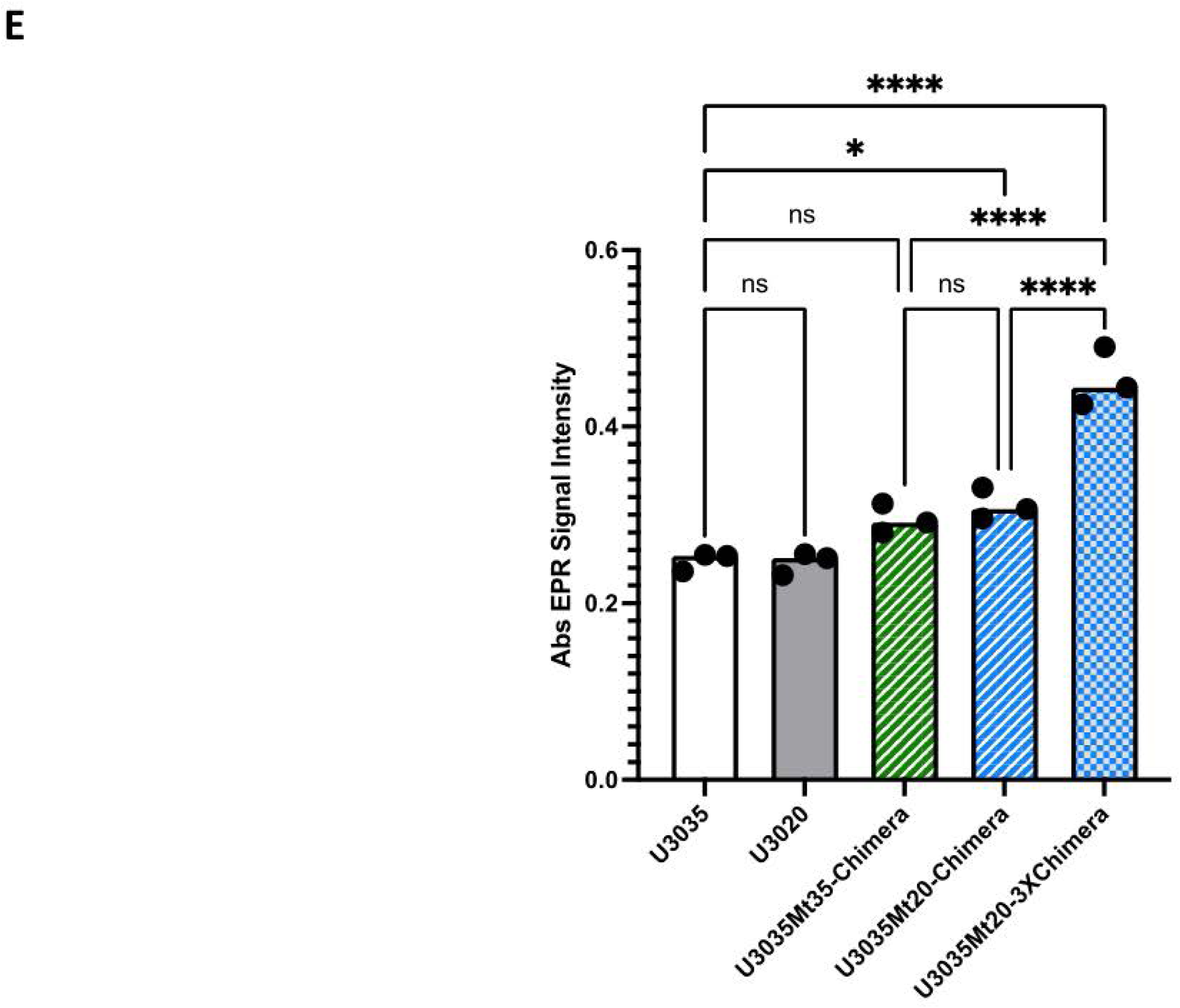
EPR spectra of DMPO adducts. **A)**. U3035 cells + DMPO (200mM). **B)**. U3035 cells + DMPO (200mM) radiated using 8Gy X-ray. **C)**. U3035 cells transplanted with mitochondria (1X) + DMPO (200mM) and radiated using 8Gy X-ray. **D)**. U3035 cells transplanted with mitochondria (3X) + DMPO (200mM) and radiated using 8Gy X-ray.

**Figure 5.**
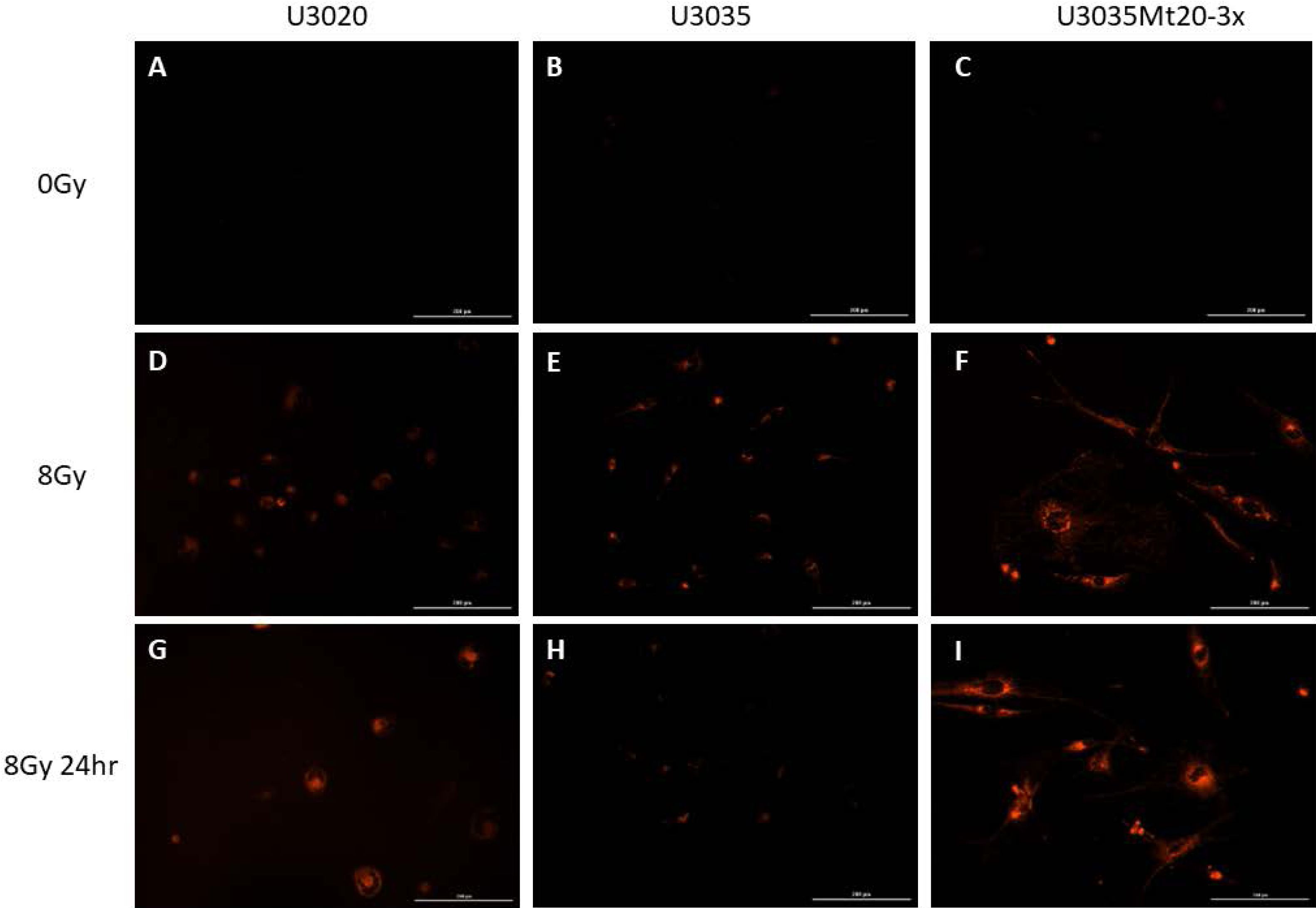
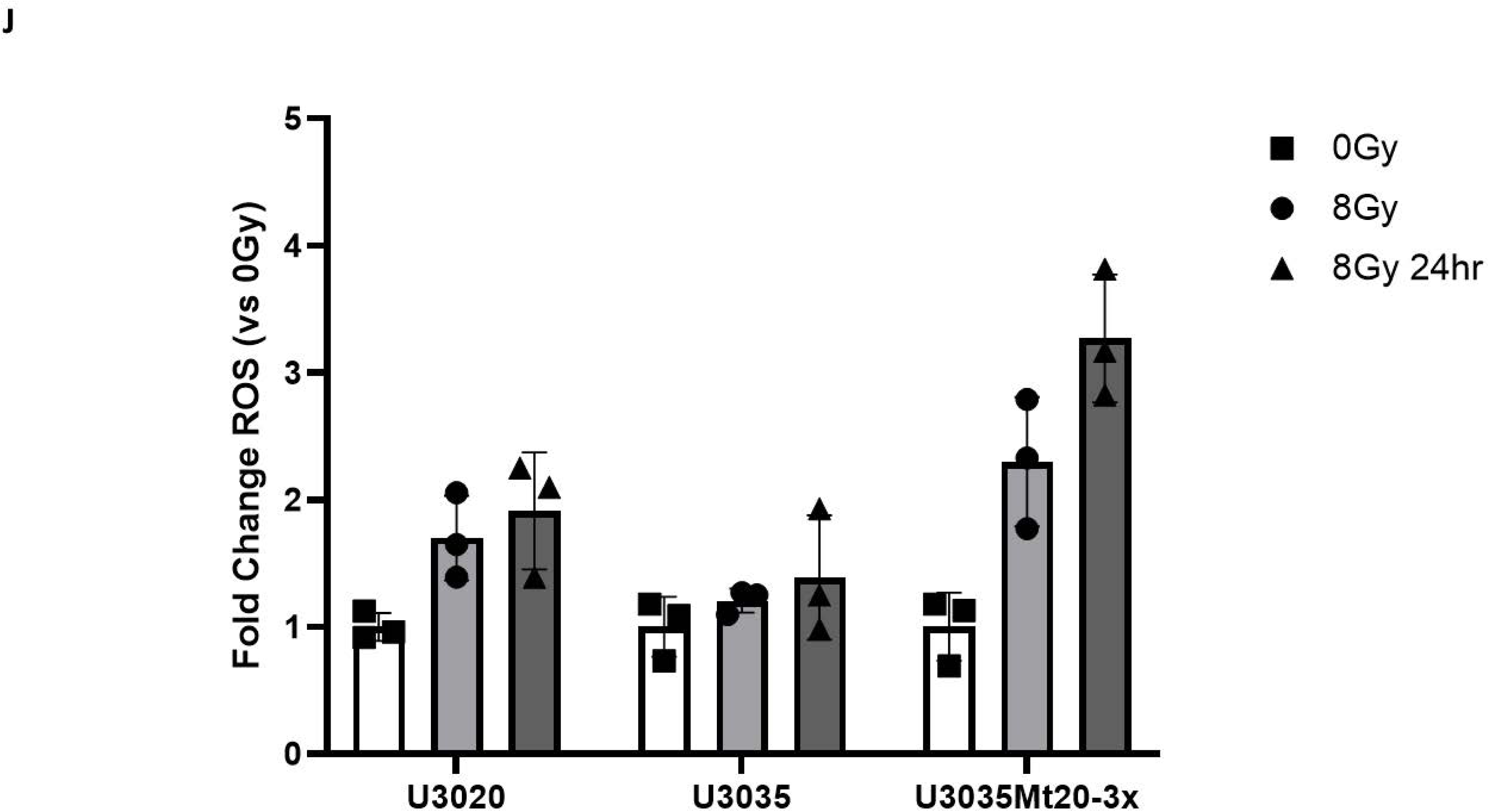
Fluorescent Probe of ROS Generation. Representative Images of: U3020, U3035, and U3035Mt20-3x cells stained with CellROX Deep red fluorescent ROS probe following 0Gy X-ray irradiation **(A-C)**. U3020, U3035, and U3035Mt20-3x cells stained with CellROX Deep red fluorescent ROS probe following 8Gy X-ray irradiation and assayed 1hr following treatment **(D-F).** U3020, U3035, and U3035Mt20-3x cells stained with CellROX Deep red fluorescent ROS probe following 8Gy X-ray irradiation and assayed 24hr following treatment **(D-F). G).** Quantification of these data represented by mean fold-change of relative fluorescent units vs 0Gy as a control.

Interestingly, when 20Gy irradiation doses were examined, there were no statistically significant differences between cell lines observed at 0.5hr post-irradiation and cytotoxicity at 24hr limited cell analyses for all groups. However, when 24hr signals were compared for 8Gy, there were significant differences observed between the GBM lines. U3020 had a sustained increase in ROS level (1.9-fold increase) while U3035 did not (1.2-fold increase). The 3.3-fold increase in the 24hr ROS signal observed in the U3035Mt20-3X chimeras was significantly greater than those observed in both U3020 and U3035 (p<0.001 for both).

### Mitochondrial transplants were durable

Analysis of mitochondrial DNA (mtDNA) sequencing from the 14-day U3035Mt20-3X chimera revealed a total of 37 heteroplasmic positions. 32 of these heteroplasmic positions were concordant with either 3020 donor or 3035 recipient sequences (**Figure 6, *gray***). The average allele frequency for 3020 donor sequences was 91.9%, while the average allele frequency for 3035 recipient sequences was 8.1%. The high degree of similarity with donor mitochondrial sequences suggests a high mitochondrial transfection and retention rate 14 days post-transfection. The remaining 5 heteroplasmic positions had *de novo* sequences that were not detected in either of the original cell lines (**Figure 6, *orange***). All *de novo* sequences had allele frequencies below 4%. A further 7 single nucleotide polymorphisms (SNPs) detected when comparing the non-chimeric 3020 donor and 3035 recipient mitochondrial sequences were completely converted (allele frequency >99%) to 3020 donor sequences in the chimeric cell line (**Figure 6, *blue***).

**Figure 6.**
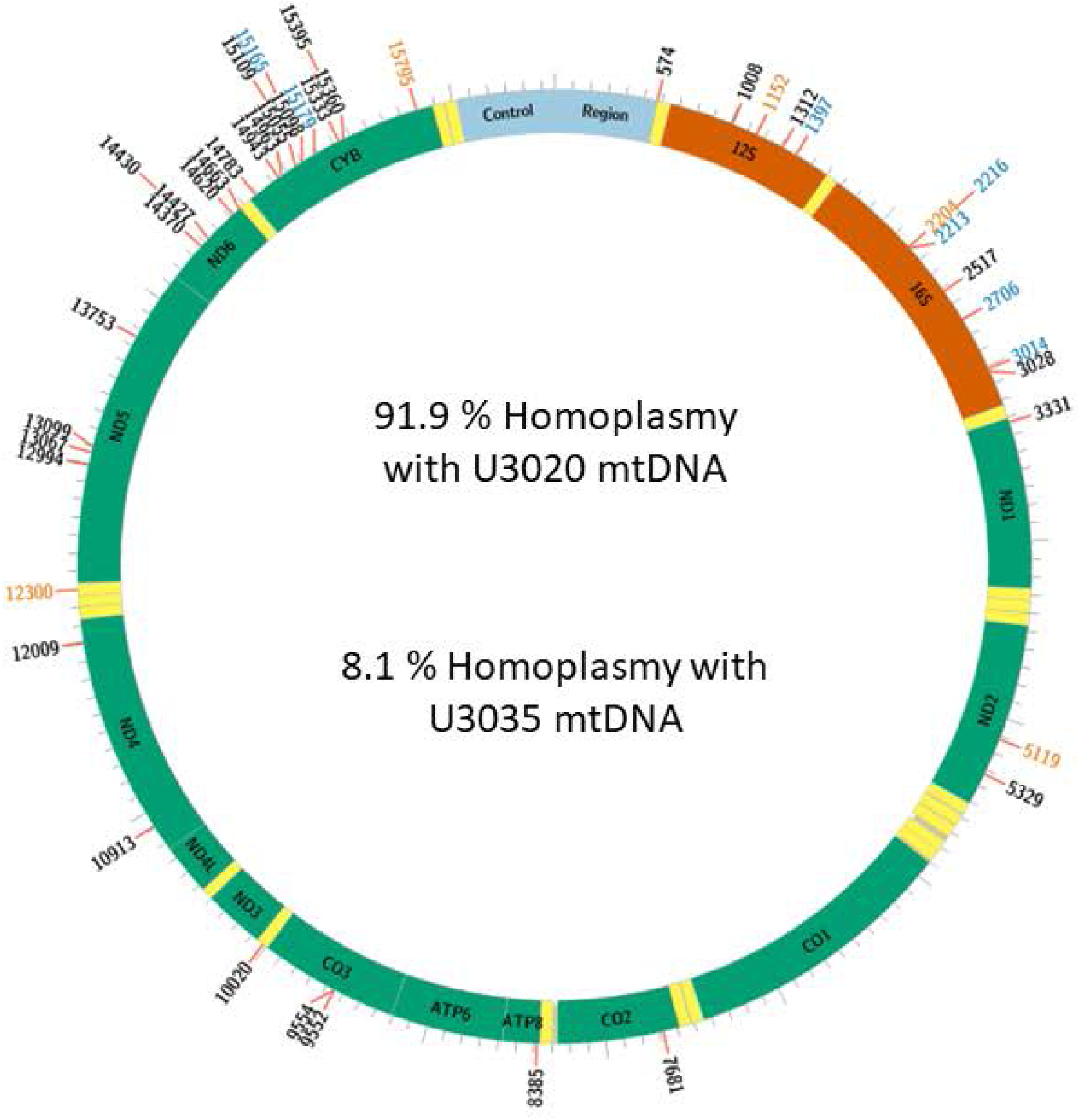
Mitochondrial DNA Sequence Analysis within U3035MT20-3x Chimeras Following Transplantation. A Circos plot designating the circular mitochondrial DNA of the U3035Mt20-x3 chimeras and the single nucleotide polymorphisms that were either shared variably with U3020 or U3035 mtDNA sequences (grey), completely converted to U3020 (blue), or not expressed in either the donor or recipient lines (orange). Allele frequencies were averaged for the grey SNP’s and combined with the blue SNPs and it was found that the DNA sequence in the 14-day post-transplant U3035Mt20-x3 chimeras was most similar to the donor U3020 line and had only retained a small proportion of the recipient mtDNA sequence.

### Mitochondrial transplant was possible in cells with an induced radio-resistant phenotype

U3035 cells were treated with 6 fractions of 2Gy Xray radiation for a total radiation dose of 12Gy to demonstrate a cell line with prior radiation exposure. Following generation of the radioresistant U3035 line, 3X allogenic mitochondrial transplants were established and persisted to post-transplant day 7 (**Figure 7**). Cells proliferated and survived past post-transplant day 10 with a subsequent loss vital dye signal.

**Figure 7.**
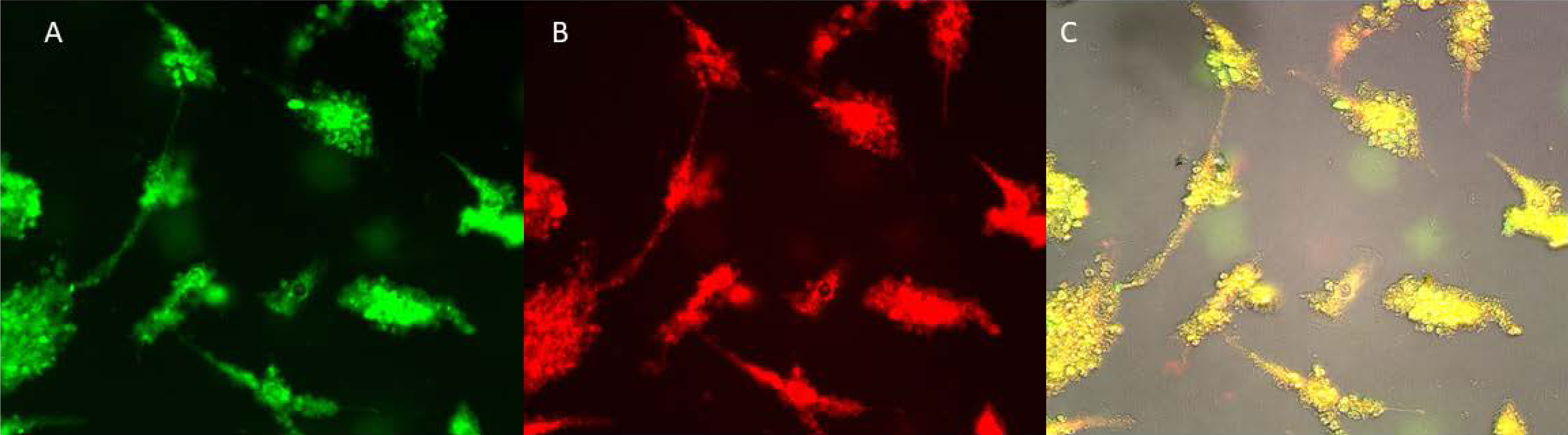
Allogenic Mitochondrial Transplantation in Radiation Resistant GBM. Representative images of **A).** Native mitochondria in radiation resistant U3035 cells stained with 200nM MitoTracker® Green vital dye. **B).** Transplanted mitochondria from U3020 cells stained with 200nM MitoTracker® Red vital dye. **C).** Overlay image of the green and red fluorescent channels highlighting successful transfer. Images were obtained at post-transplant at 48hrs.

## Discussion

Our understanding of the role of metabolic pathways in the tumorigenesis and development of treatment resistance in glioblastoma (GBM) continues to expand. The importance of isocitrate dehydrogenase (IDH) isotype mutants in the latest WHO classification system underscores the fundamental importance of tumor cell metabolism in prognostication, while IDH-based therapy offers therapeutic promise in IDH-mutant tumors [21, 22]. Although the majority of clinical interest has been placed on determining mutation/epigenetic status in diagnostic patient specimens, a wealth of preclinical data have emerged regarding the role that mitochondria play and their potential to form relevant reactive oxygen species (ROS). For example, the increased action of mitochondrial superoxide dismutase 2 (SOD2) in glioblastoma has been attributed with a role in the development of resistance to temozolomide, the first-line standard of care chemotherapeutic [23, 24]. Similarly, evidence supports the upregulation and stabilization of free radical scavenger systems within glioma stem-like cells (GSCs), thereby limiting the accumulation of mitochondrial ROS and conferring radiation resistance [25]. Based on the data in the current study, we propose a methodology for using exogenous donor mitochondria as a means by which to increase mitochondria reactive oxygen species (mtROS) in GBM cells *in vitro* in response to ionizing radiation exposure.

While the concept of mitochondrial transfer or transplantation is not new, the application of such a process to sensitize a tumor cell to standard therapy, such as radiation, is novel in the context of glioblastoma. Spontaneous mitochondrial transfer has been documented in the CNS, through which non-tumor astrocytic mitochondria transferred to GBM cells within the tumor microenvironment [26].

The presence of tunneling nanotubes (TNTs) has been demonstrated in GBM resection specimens while mitochondrial transfer has been demonstrated *in vitro* between GBM cells and adjacent non-neoplastic astrocytes, conferring resistance to hypoxia and increased glucose- and glutamine-dependence [27]. In the current study, we enlisted the aid of the cell-penetrating peptide (PEP-1; sequence KETWWETWWTEWSQP-KKKRKV) as a means of artificially introducing both allogeneic and autogenic mitochondria into the U3035 GBM cell line, previously shown to exhibit a rapid decay in ROS generated in response to radiation exposure, likely due to effective scavenger systems [28]. In doing so, we were successful in confirming that relative mitochondrial content does correlate to an increase in ROS, detectable via EPR spin-trapping techniques and mitochondrial dye tracking. These findings are concordant with those in other cell types, namely skeletal muscle [29].

The sustainability of donor mitochondria post-transplantation was also of interest in developing a methodology that would eventually have a practical therapeutic potential use. Whereas the native U3035 GBM cells demonstrated a rapid decay of ROS both in EPR and mitochondrial staining assays, the U3020 GBM cells maintained mitochondrial ROX-staining for at least 24 hours post-radiation. With ROS longevity typically limited to milliseconds, thus requiring spin-trapping solvents to confirm their presence, to maintain a durable ROS signal via mitochondrial staining is a promising finding. The durability of the transplantation was also confirmed using mtDNA sequencing, indicating that the donor mitochondrial are retained in the recipient cell for at least two weeks, well beyond the standard doubling time of the U3035 GBM cells. Thus, the CPP-mediated mitochondrial transplantation from one GBM cell line to another appears to be achievable and durable within the context of our time course.

One feature that complicates the development of novel therapeutics in GBM research is that nearly all clinical trials are initially performed in the setting of recurrent disease after the delivery of standard of care first-line chemotherapy and external beam radiotherapy. Hence, we endeavored to explore the feasibility of using the CPP mitochondrial transplantation on GBM cells that were already radiation resistant. In generating U3035 cells exposed to serial radiotherapy doses, we confirmed that the CPP methodology can deliver donor mitochondria to radiation resistant cells, as would be the likely scenario in clinical investigation. Further studies will be needed to confirm if the ROS generated in response to subsequent radiation post-transplantation is similar to the radiation naïve cell studies.

The contribution of ROS as a tumorigenic versus therapeutic agent in setting of malignancy remains the subject of debate [30]. Whereas basal increases in ROS levels have been associated with neoplastic transformation, breaching the threshold past which ROS levels become toxic within tumor cells appears to be a viable therapeutic strategy as demonstrated by the numerous cancer treatments tasked with that same goal [31]. The fundamental question remaining to be answered pertains to the identification of the level at which ROS becomes lethal and if that level can be achieved using mitochondrial harvested from autologous tumor cells or autologous non-neoplastic tissue. As previously discussed, the stoichiometry of ROS production based on mitochondrial content in the present study mimics that of work completed in skeletal muscle [29]. From a practical perspective, one future area of study would be to examine the ability to successfully transplant mitochondria from skeletal muscle for determination of both ROS potential and durability. In the setting of recurrent GBM undergoing surgical resection, options for subsequent radiation delivery are very limited, usually using hypofractionated or radiosurgery techniques [32, 33]. During surgery, local muscle is usually accessible as part of the standard approaches. Harvesting a small amount of mitochondria-dense tissue with CPP preparation and injection into the presumption post-operative radiation target areas could afford a novel mechanism for increased lethal ROS generation.

Although we are able to conclude that our technique of mitochondrial transplantation into GBM cells was successful based on ROS generation and durability endpoints, there are limitations to the work. First, in the attempt to provide data on human-derived tumor cells, no *in vivo* rodent studies have been performed. With respect to the EPR data, the availability of effective *in vivo* ROS probes is extremely limited. Moreover, the proposed application of use in recurrent GBM, creates a further obstacle to create an immune-competent rodent model where data beyond the tolerability of surgery, radiation, and CPP use could be tested. Perhaps the greatest limitation of the current study lies in the lack of predictability of ROS lethality and the level to which intracellular ROS needs to reach in order to insure radiation-induced cell death. Further studies that incorporate the concurrent blockade of free-radical scavenger systems will be needed to optimize the techniques in order to achieve maximal results while minimizing radiation exposure.

## Materials and Methods

### Cell Lines

Two human mesenchymal subtype glioblastoma cell lines, U3035 and U3020, were obtained from the Human Glioblastoma Cell Culture (HGCC) Biobank at Uppsala University in Sweden. The cells were cultured in defined media (DMEM/F-12, EGF, FGF, N2, B27 supplements) with penicillin/streptomycin at 37°C in 5% CO_2_ according to recommended passaging parameters.

Verification of cell line authenticity was performed via short tandem repeat (STR) analysis by ATCC in Manassas, VA, USA (*supplemental data*). A subset of U3020 and U3035 cells were cultured under hypoxic conditions (1% O_2_) to assess native mitochondrial response to low oxygen levels using a C-Chamber Cytocentric® incubator subchamber (Biospherix, Parish, NY, USA).

### Mitochondrial Isolation and Quantification

U3020 and U3035 cells were cultured in T75 flasks until 90% confluency was achieved. Cells were dissociated with Accutase® (Milipore Grp, Burlington, MA), pelleted via centrifugation, and resuspended in 1mL mitochondrial isolation buffer. Cell counts were obtained using a Cellometer® Auto 2000 cell viability counter (Nexcelom™ Bioscience LLC, Lawrence, MA). Mitochondria were then isolated using a commercially available kit according to manufacturer’s instructions (Biovision™, Milpitas, CA, USA). Briefly, reagent-based permeabilization was performed on whole cells to allow for mitochondrial extraction from the cytosolic space. Differential centrifugation first separated mitochondria from other cellular debris (600xg) and then isolated mitochondria from other biomolecular components of the cytosolic space (7000xg). Mitochondria were resuspended in a storage buffer and the total protein content of the mitochondria isolates were measured using a TAKE-3 plate (Bio Tek; Charlotte, VT) and optical density A280 quantification measurement.

### Mitochondrial Labeling and Transplantation

Whole cells or mitochondria were labeled using Mitotracker® (ThermoFisher™) vital dyes. Recipient U3035 cells were seeded at a density of 5.0×10^5^ cells/well of 12-well Corning CoStar geometry plate and allowed to settle overnight. Media in each well was replaced with serum-free neural basal media containing 200nM green MitoTracker® dye and allowed to incubate at 37°C 5.0% CO_2_ for 30 minutes. Donor mitochondria were isolated as described above and resuspended in neural basal media containing either 200nM green (for post-transplant quantification) or red (proof-of-concept) MitoTracker® dye for 30 minutes at 37°C 5.0% CO_2_. Dyed mitochondria were then incubated with a cell-penetrating peptide, Pep-1-Cysteamine (Pep-1) (suspended in 1X phosphate-buffered saline), at a ratio of 0.06mg Pep-1 / 100µg of mitochondrial protein in 400µL of PBS, as described by Chang, et al [34]. The mitochondria and Pep-1 were allowed to conjugate for 20 minutes at room temperature. Following the Pep-1 conjugation, either 400µL of 100µg of mitochondrial protein (1X transplant) or 300µg of mitochondrial protein (3X transplant) were added to U3035 cells cultured in 12-well plates containing DMEM. Mitochondria were allowed to transplant over a period of 48hr after which fresh DMEM was replaced. Images of successful transplants were taken at 48hr and 168hr (7d) on an ECHO inverted microscope (BICO; Gothenburg, Sweden) with FITC and TRITC channels. Resulting mitochondrial chimeras were continued in culture for 14 days to determine transplant durability. Fluorescent intensity of 3X chimeras were quantified using ImageJ software [35].

### Mitochondrial DNA Extraction

Mitochondrial DNA was extracted directly from freshly isolated mitochondria following the protocols outlined by Abcam (Cambridge, United Kingdom). Briefly, an enzymatic mixture (proteinase K and RNAse) was allowed to incubate with the mitochondria for 30 minutes at 55°C. Mitochondrial samples were then mixed with an alkaline lysis buffer (containing NaOH), allowed to incubate on ice for 5 minutes. An acidic neutralization buffer (containing glacial acetic acid) was then added to the sample tubes. Mitochondrial DNA samples were then separated via centrifugation and the supernatants were added to DNA trapping spin columns. This supernatant was spun through the columns and washed twice using a wash buffer containing 75% ethanol. DNA samples were then eluted with a Tris-HCl/EDTA buffer at room temperature for 2 minutes. Samples were recovered from columns using centrifugation and DNA quality and quantity were measured using A_260_ determinations through optical density methods and the Take3 Bio Tek platform.

### Mitochondria DNA Sequencing and Analyses

Mitochondrial DNA concentration were quantified using a Qubit 4 Fluorometer (ThermoFisher Scientific, Waltham, MA). The SparQ DNA Fragment and Library Prep kit was used to build the mitochondrial DNA libraries using 1 to 4 ng of template according to protocol. The completed libraries were quantified using the Qubit and run on an Agilent Bioanalyzer with the High Sensitivity DNA chip to determine the average size of each library. The libraries were pooled together, at an equal molar ratio, and run on a Miseq with an Illumina V2 300 cycle (PE150bp) reagent kit.

Adapter sequences were trimmed from all FASTQ files using Atria (version 4.0.3; RRID:SCR_021313). Successful removal of adapter sequences was confirmed using FastQC (version 0.12.1; RRID:SCR_014583). NOVOPlasty (version 4.3.5; RRID:SCR_017335) was then used for *de novo* assembly of the mitochondrial genome using adapter trimmed deep sequencing reads from the U3020 parental line with the following input parameters: genome size constraint 12-22 kilobases, K-mer of 33, and DNA sequence for MT-ATP8 (gene ID: 4509) from GRCh38.p14 as the seed. Single nucleotide polymorphisms (SNPs) in the U3035 parental line relative to the constructed U3020 mitochondrial genome assembly were detected using the heteroplasmy function of NOVOPlasty. A total of 39 such SNPs were detected at homoplasmic allele frequencies (≥0.99) in the U3035 parental line.

The U3035 14-day U3035Mt20-3X chimera was then compared to the same U3020 mitochondrial genome assembly. A total of 37 SNPs were detected at heteroplasmic allele frequencies (0.01-0.99) with 32 of the 37-sharing identity with homoplasmic SNPs identified in the U3035 parental line. Percent similarity was then determined by averaging the variant allele frequencies for the 32 shared SNPs. All data were graphically presented using Circos circular ideogram visualization [36].

### Cell Irradiation and Reactive Oxygen Species Measurements

U3035, U3020, U3035 autologous chimeras (U3035Mt35-Chimeras), U3035 allogenic chimeras (U3035Mt20-Chimeras), and U3035 allogenic 3X chimeras (U3035Mt20-3XChimeras) were exposed to 8Gy of ionizing radiation (220kV, 13mA) using a XenX radiation platform (Xstrahl; Suwanee, GA) to determine if there were differential generation of reactive oxygen species (ROS) between the cell lines. 5.0×10^5^ cells were suspended in 200mM DMSO, irradiated, and subsequently probed using electron paramagnetic resonance (EPR) via and ELEXSYSII spectrometer (Bruker; Billerica, MA) for quantification of hydroxyl radical production. All lines were also probed for non-specific ROS generation using CellROX® red dyes (ThermoFisher Scientific; Waltham, MA). Briefly, cells were given 5nM of the CellROX® dye in their native 6-well plate and DMEM immediately prior to exposure to 8 or 20Gy of ionizing radiation (220kV, 13mA). Lines were assayed again 24hr following irradiation. Non-irradiated, CellROX® treated cells and irradiated cells with no CellROX® reagent were used as positive and negative controls, respectively. Samples were allowed to incubate at 37°C 5.0%CO_2_ for 30 minutes. TRITC images were obtained for all experimental conditions and fluorescent intensity was quantified using ImageJ software. A subset of U3035 cells were irradiated at 2Gy (220kV, 13mA) every other day, for 12 days, in order to generate a 12Gy radioresistant cell line to determine if mitochondrial transplantation was feasible in cells which are not radiation naïve.

## Data availability statement

The datasets generated and analyzed during the current study are available in the NCBI BioProject repository under accession number PRJNA1134145 or individual sequences at NCBI Sequence Read Archive (SRA) under accession numbers SRX25532763, SRX25532762, and SRX25532761.

## Author contributions

CPC and KLM made substantial contributions to the design of the work, data acquisition, analysis, and drafting of the manuscript. MV, VVK and AM contributed to data acquisition and manuscript preparation. All authors reviewed final draft prior to submission.

